# Controlled Release of Poly(U) via Acetalated Dextran Microparticles for Enhanced Vaccine Adjuvant Delivery

**DOI:** 10.1101/2025.07.16.665134

**Authors:** Sophia A. Ly, Nicole Rose Lukesh, Erik S. Pena, Ryan N. Woodring, Sophie E. Mendell, Connor T. Murphy, Grace L. Williamson, Kierstin A. Clark, Alexandra M. Lopez, Eric M. Bachelder, Kristy M. Ainslie

**Author notes:** Corresponding author: Kristy M. Ainslie, Fred Eshelman Distinguished Professor, Division of Pharmacoengineering & Molecular Pharmaceutics, UNC Eshelman School of Pharmacy, 4012 Marsico Hall, 125 Mason Farm Road, Chapel Hill, NC 27599, United States.

## Abstract

Diverse drug delivery systems are needed to address challenges in delivering novel vaccine components and enhancing their efficacy. Poly(U) is a single-stranded RNA composed of uracil repeats that acts as a toll-like receptor (TLR) 7/8 agonist, stimulating the innate immune system. However, poly(U) is susceptible to ribonuclease degradation without a delivery carrier, and its negative charge hinders cellular uptake. Encapsulation in acetalated dextran (Ace-DEX), a pH-sensitive, biodegradable polymer, addresses these challenges. This study encapsulated poly(U) into Ace-DEX Microparticles (MPs) with either spherical (smooth MPs) or collapsed-surface (wrinkled MPs) via spray-drying. It was hypothesized that the different morphologies of MPs would influence the vaccine efficacy after in vitro and in vivo models. Smooth poly(U) MPs had a higher percent viability and cytokine response in dendritic cells (DCs) than wrinkled poly(U) MPs. Moreover, mice vaccinated with smooth poly(U) MPs + ovalbumin (OVA) showed enhanced IL-2 production and IFN-γ in response to OVA peptide and MHC-I immunodominant peptide restimulation, respectively, compared to wrinkled poly(U) MPs. However, mice vaccinated with wrinkled poly(U) MPs + OVA significantly increased B-cell and germinal center B-cell frequencies compared to mice vaccinated with phosphate buffered saline (PBS) whereas mice vaccinated with smooth poly(U) MPs + OVA did not. Overall, these findings suggest that smooth poly(U) MPs modulated dendritic cells and T-cells, and wrinkled poly(U) MPs modulated B-cells. Understanding how morphology influences these cell types will aid in optimizing future vaccine systems for more specific cellular targeting.

## Introduction

Worldwide, vaccines are estimated to prevent four million deaths each year [1]. Vaccines prime the immune system to specifically recognize and mount an immune response against pathogens. Among these, subunit vaccines represent a promising approach. Subunit vaccines can be composed of selected protein antigens specific towards a pathogen for a targeted immune response [2]. This precision minimizes the risk of adverse effects commonly associated with live virus vaccines. However, a challenge of subunit vaccines is their inherent low immunogenicity in that the protein antigens alone often fail to stimulate a protective immune response [3]. To overcome this limitation, adjuvants are included in vaccine formulations to enhance the immune response to the vaccine antigens, improving the vaccine’s efficacy [4].

One such adjuvant is poly(U), a single stranded RNA (ssRNA) that acts as a toll-like receptor 7/8 (TLR7/8) agonist. TLRs are pattern recognition receptors (PRRs) that respond to pathogen-associated molecular patterns (PAMPs) and initiate immune responses [5, 6]. By binding TLR7/8, poly(U) activates antigen-presenting cells (APCs) and enhances immune responses [7, 8]. A similar adjuvant that has been studied is poly(A:U). Poly(A:U) is a TLR 3/7 agonist that is composed of poly(A) and poly(U). Poly(A:U) increases cytokine production in bone marrow dendritic cells and in vivo [9]. (Poly(A:U) has also improved the delivery of the cancer drugs 5-fluorouracil and adriamycin in patients with operable gastric cancer [10]. While previous studies found that poly(A) and poly(U) had little enhancing effects on immune responses, studies that formulated poly(A) into polymeric nanoparticles that were immunologically impactful [11, 12]

The results from delivering poly(A) through nanoparticles implores the exploration of poly(U) as a promising adjuvant for enhancing the efficacy of subunit vaccines by allowing for rapid immune system responses and generation of long-lived memory cells. However, the previous inefficacy may be because TLR7/8 are located intracellularly in the phagosome and due to its negative charge, poly(U) has difficulty crossing the cell membrane to bind to its intracellular cognitive pattern recognition receptors [5, 13]. This necessitates a drug delivery carrier capable of cellular internalization to make a robust vaccine.

Traditional drug delivery carriers such as polylactic-co-glycolic acid (PLGA) can reduce the local pH to cause toxicity, and inflammation through its acid byproducts [14-16]. Additionally, even though PLGA degradation rates are tunable (dependent on lactic acid and glycolic acid ratios), it still takes weeks to months to degrade, which can limit its use [17]. mRNA based lipid nanoparticles (LNPs) are another common drug delivery carrier; however, their structures may cause them to have poor drug loading efficiency for a charged molecule like poly(U) [18]. Additionally, it is imperative to keep vaccines affordable, and therefore production costs must stay low. The World Health Organization stopped developing a PLGA-based tetanus vaccine partly because it was too expensive and challenging to manufacture at scale [19]. The PLGA vaccine relied on batch formulation through coacervation, rather than more scalable techniques like spray drying. In comparison to LNP, protein-based vaccines are less expensive and faster to make than mRNA vaccines. Depending on the cell system used (e.g. insect, plant, or yeast), 10 million doses of a protein vaccine can be made in 30 days for about $0.001 per dose. On the other hand, just the mRNA in an mRNA vaccine can cost over $1 per dose [20-23]. Therefore, new formulation approaches need to be considered for the application of a vaccine adjuvant like poly(U).

Acetalated dextran (Ace-DEX) is an acid-sensitive polymer with tunable degradation rates dictated by cyclic acetal coverage (CAC). The polymer is synthesized by reacting dextran with 2-ethoxypropene, introducing both cyclic and acyclic acetal groups whose ratio determines degradation rate [24]. There are many methods to formulate Ace-DEX MPs to encapsulate poly(U), however spray drying is advantageous because of its scalability and alignment with GMP industry standards [25]. Spray drying is a one-step, continuous process that consists of spraying a drug-polymer solution through a pressure nozzle into a heated chamber [26]. This process forms Ace-DEX MPs with ∼1 µm diameter. This is advantageous because 1–2 µm particles were taken up about 50 times more by dendritic cells (CD11c^+^) than by macrophages (CD11c^−^), suggesting that larger particles are primarily transported to lymph nodes by dendritic cells [27]. Therefore, Ace-DEX MPs passively target phagocytic APCs based on their size and once internalized they readily degrade in the acidic environment of the cell’s phagosome releasing the adjuvant where TLR 7/8 resides. [5, 24]

MP morphology can influence how readily they are phagocytosed. Spherical MPs are more rapidly internalized by cells than rod-shaped MPs [28-30]. However, other studies have found that MPs with angular surfaces can have even faster internalization compared to smooth, round MPs. The shapes of the MPs impact how they adherence to the cell membrane and actin filaments for endocytosis [31]. Our previous work has used a variety of MP formulations with different morphologies for vaccines in autoimmunity and viral infections [32-34]. Here we report how spray drying conditions and microparticle morphology, specifically smooth versus wrinkled Ace-DEX MP, influence vaccine performance, based on the hypothesis that encapsulating poly(U) in these particles enhances activation of antigen presenting cells and promotes a proinflammatory immune response compared to soluble poly(U).

## Materials and Methods

### Materials

All chemicals were purchased through Sigma Aldrich (St. Louis, MO) unless stated otherwise. All biologics, assays, and disposables were purchased from Thermo Fisher Scientific (Waltham, MA) unless stated otherwise. GraphPad Prism 10 was used to generate, format figures and perform ANOVA analysis.

### Ace-Dex Synthesis

Synthesis of Ace-DEX was performed with 71 k dextran from *Leuconostoc mesenteroides* in dimethyl sulfoxide (DMSO) [35]. The catalyst, pyridinium *p*-toluenesulfonate and 2-ethoxypropene were then added to the mixture to begin the reaction to create Ace-DEX in anhydrous conditions. After fifteen minutes, tetrahydrofuran (THF) was added and two hours later triethylamine (TEA) was added to quench the reaction. The Ace-DEX was precipitated in basic water (0.04% TEA by volume in water), isolated by centrifugation, and lyophilized overnight. For further purification, the lyophilized product was dissolved in ethanol and centrifuged. The supernatant was precipitated in basic water and lyophilized to yield Ace-DEX polymer. To characterize the polymer, ^1^H 400 MHz NMR (Inova) was used to determine the cyclic acetal coverage (CAC) of the Ace-DEX (60% CAC).

### Microparticle Fabrication

MPs were formed via spray drying using Mini Spray Dryer B-290 (Büchi, New Castle, DE). The wrinkled MPs were dissolved in a 90:10 ethanol:PBS solvent system with poly(U). The smooth MPs were dissolved in 99:100 ethanol:DMSO solvent system. All MPs were spray dried with parameters listed in Table 1. The MPs were then dissolved in a 20 mg/mL solution of sucrose and basic water, frozen, and lyophilized.

**Table 1.** Spray Drying Parameters.

|  | WRINKLED MPS | SPHERICAL MPS |
| --- | --- | --- |
| LOADING (%) | 1.0 | 1.0 |
| CONCENTRATION (MG/ML) | 5 | 2.5 |
| **INLET FLOW (RPM)** | 28 | 28 |
| **Q FLOW (RPM)** | 60 | 60 |
| **TEMPERATURE (°C)** | 75 | 75 |
| **SOLVENT SYSTEM** | 90% EtOH:10% H<sub>2</sub>O | 99% EtOH:1% DMSO |

**Table 2.** MP characterization of smooth and wrinkled MPs showing EE of poly(U) and MP diameter (nm).

|  | WRINKLED MPS | SMOOTH MPS |
| --- | --- | --- |
| DRUG LOADING (MG/ML) | 1 | 1 |
| DIAMETER (NM) | $1300 \pm 698$ | $812 \pm 190$ |
| ENDOTOXIN (EU/ML) | $0.0598 \pm 0.0099$ | $0.0103 \pm 0.0548$ |

### Microparticle Characterization

To characterize the diameter, MPs were suspended in basic water and analyzed by dynamic light scattering (Brookhaven NanoBrook 90Plus Zeta Particle Size Analyzer, Holtsville, NY). For scanning electron micrographs (SEM) (Hitachis-4300 Cold Field Emission SEM; Ibaraki, Japan; UNC CHANL), MPs were suspended in basic water and a droplet was deposited onto carbon tape on top of an SEM stub and dried overnight. To screen for endotoxin, all MPs were tested using the Pierce LAL Chromagenic Endotoxin Quantification kit. All MPs were reported to be below the 0.1 EU/mg endotoxic concentration. The encapsulation efficiency (EE) of the MPs was completed using nanodrop by dissolved MPs in ethanol (Thermo Scientific NanoDrop One Microvolume UV-Vis Spectrophotometer).

### Release Study

To observe particle degradation, MPs were suspected in phosphate buffered saline (PBS, pH 7.4) at 1 mg/mL. The suspended MPs were placed in a 5 mL Eppendorf tube and placed on a shaker plate set to 37 °C to mimic physiological conditions. 300 µl aliquots of the suspended MPs were taken at several time points. The aliquot was then centrifuged to pellet the MPs. The supernatant was then removed and the pellets were stored in 20 °C freezer. Once pellets from each timepoint were collected and processed, they were prepared, and the amount of poly(U) was measured through nanodrop.

### Dendritic Cell Stimulation

DC2.4 cells in Roswell Park Memorial Institute (RPMI) 1640 media (with 10% by volume fetal bovine serum and 1% by volume of penicillin-streptomycin) were seeded at 25,000 cells/well in a 96 well plate and incubated overnight. MPs were suspended in media and added to the cells at a poly(U) concentration of 13.5, 27.0, 54.0, 108.0, and 215.0 ng/mL and incubated for 24 hours. Blank MPs were added at corresponding MP concentration. Untreated controls were also included. Supernatant was then taken from the cells to measure cell cytotoxicity by lactose dehydrogenase (LDH) (Invitrogen, San Diego, CA), cell viability by Cell Titer Blue, and cytokines by sandwich ELISA (Biolegend, San Diego, CA).

### In vivo vaccination

Mouse studies followed the guidelines set by the National Institute of Health and were approved by the UNC Institutional Animal Care and Use Committee (IACUC). Female BALB/c mice (n = 5 per group) that were 8-10 weeks old from Jackson Laboratories (Bar Harbor, ME) were vaccinated subcutaneously on days 0, 21, and 35 with 10 µg of poly(U) (InvivoGen) and 10 µg of OVA.

### Antigen Recall Experiments

Splenocytes and inguinal lymph node lymphocytes were isolated from mice vaccinated with OVA and different MP formulations. Cells were plated in 96-well U-bottom plates (5×10^5^ cells/well) and stimulated with 10µg/mL of OVA protein, MHCI OVA peptide (SIINFEKL), or MHCII OVA peptide (ISQAVHAAHAEINEAGR) for 36hr. Cells were also used to assess antigen-specific IFN-γ and IL-2 via ELISpot (BD Biosciences; San Jose, CA). Unstimulated samples from each mouse were prepared and their cytokine levels or spots were subtracted from stimulated samples.

### Flow Cytometry Characterization

Splenocytes and inguinal lymph node lymphocytes were stimulated for 36hr as above and stained for flow cytometry using the following panel: eBioscience Fixable Viability Dye eFluor 506 (diluted 1:1000) and the following fluorescent antibodies from Biolegend (San Diego, CA): CD3 Alexa Fluor 488 (diluted 1:100), CD4 APC/Fire 750 (diluted 1:200), CD8 PerCP/Cy5.5 (diluted 1:100), CD44 Brilliant Violet 421 (diluted 1:400), CD62L Brilliant Violet 785 (diluted 1:400), CD19 APC (diluted 1:400), GL7 PE (diluted 1:1600), and CD38 PE/Cyanine7 (diluted 1:800) **(Fig S1, Fig S2)**.

## Results and Discussion

There is a limited number of FDA approved adjuvants for vaccines, and new ones are needed. Since poly(U) has difficulty crossing the cell membrane without a drug delivery vehicle, it was encapsulated in Ace-DEX MPs by spray drying. Previous studies have demonstrated that the morphology of drug delivery carriers can influence cellular uptake and release kinetics [28-30]. Spherical particles have been found to be readily taken up intercellularly [36]. Additionally, other studies have shown that MPs with angular surfaces can be more quickly internalized than smooth, round MPs due to geometric adherence to cells [31]. To investigate this, we used two solvent systems to generate MPs with either wrinkled or smooth morphologies. Spray drying with ethanol and water produced wrinkled particles due to the relatively low boiling points of these solvents (78 and 100°C, respectively). In contrast, spray drying with ethanol and DMSO yielded smooth particles. The higher boiling point of DMSO (189°C) slows solvent evaporation during spray drying, promoting a smoother, spherical morphology. To characterize Ace-DEX MPs, the encapsulation efficiency (EE) of poly(U) was quantified. Smooth MPs had a slightly higher(?) EE of 43.20 ± 3.20 % EE, whereas wrinkled MPs had an EE of 28.87 ± 4.55 % EE. Smooth MPs had a radius of 812.1± 189.8 nm and wrinkled MPs were measured at 1300.1 ± 698.6 nm. These differences in EE and diameter were statistically significant.

Successful adjuvants have controlled release to sustain and strengthen the immune response. To assess how each MP retained poly(U), release profiles were observed by suspending MPs in PBS at 37° to mimic physiological conditions. Both particle sets had observed burst release where approximately 40% of the encapsulated cargo was released at the first time point. Over the first six hours, this percentage increased to approximately 80% **(Fig 1B)**, which gradually increased to approximately 95% release by day 7 **(Fig 1C)**. Though the smooth poly(U) MPs exhibit increased release, particularly around the 6 hour timepoint, both MP morphologies have similar release profiles.

**Figure 1.**
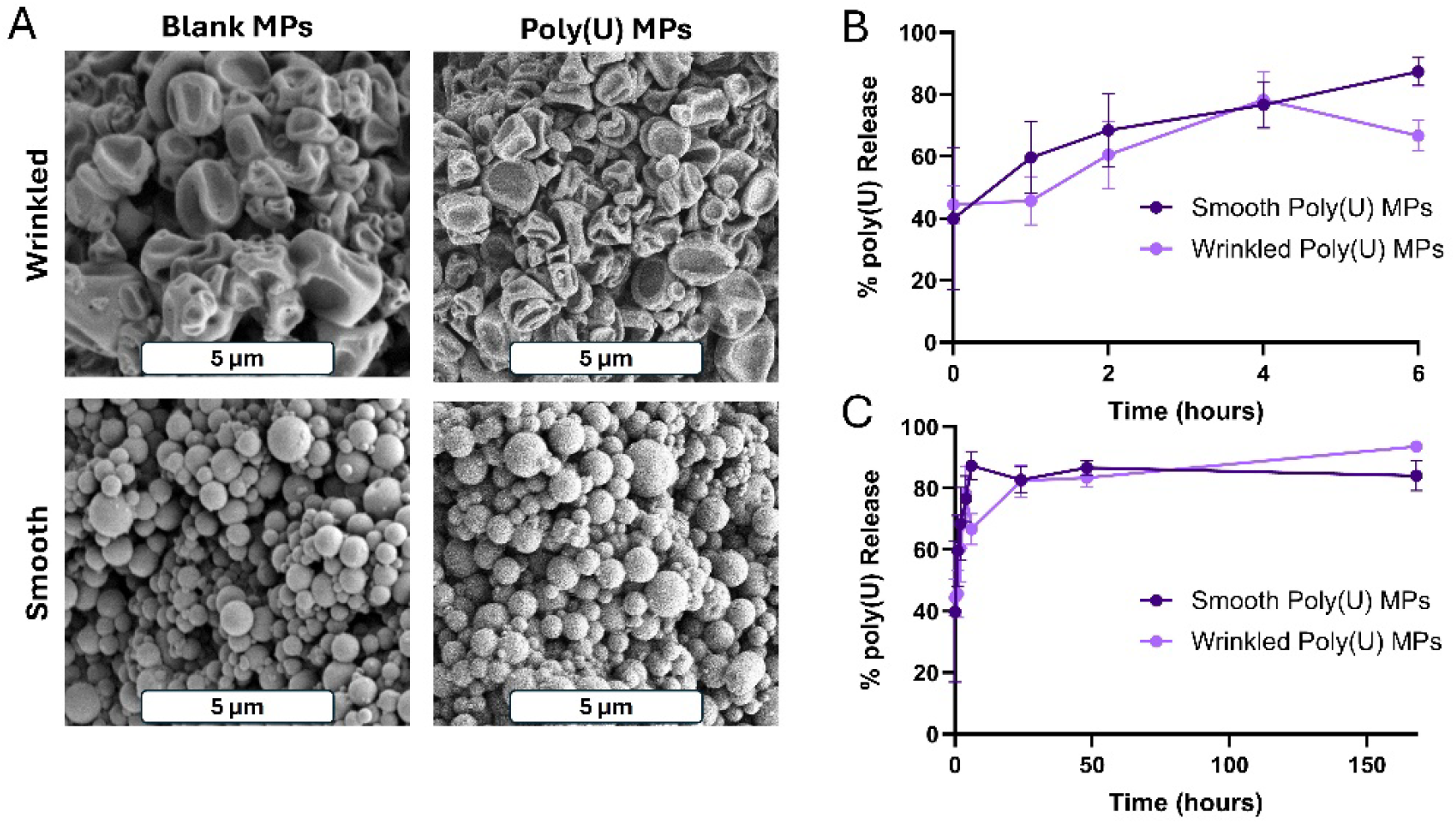
Spherical and wrinkled poly(U) MPs formulated by spray drying exhibit similar release profiles. Scanning electron microscopy images of (A) wrinkled blank MPs, poly(U) encapsulated wrinkled MPs, smooth blank MPs, poly(U) encapsulated smooth MPs. Poly(U) release profiles over (B) first 6 hours and over (C) one week. Scale bar is 5µm. Data presented as mean ± standard deviation.

We have previously shown that Ace-DEX MPs are readily taken up by dendritic cells (DCs) and enhance antigen cross presentation compared to other formulations [37]. DCs are critical for the immune response as they educate adaptive immune cells such as B cells and T cells. Therefore, to understand if poly(U) MPs of different morphologies activate DCs, DC2.4s were incubated with poly(U) MPs and soluble poly(U) for 24 hours. Viability, cytotoxicity, and cytokines were measured. Only at the highest poly(U) concentration of 215 ng/mL were significant differences noted in viability between smooth and wrinkled poly(U) MP groups, where smooth poly(U) MPs had a higher viability than the wrinkled poly(U) MPs **(Fig 2A)**. CellTiter Blue results were confirmed by LDH **(Fig S3A)**. Utilizing both methods to measure cell health provides information about metabolic activity of the cells as well as membrane integrity.

**Figure 2.**
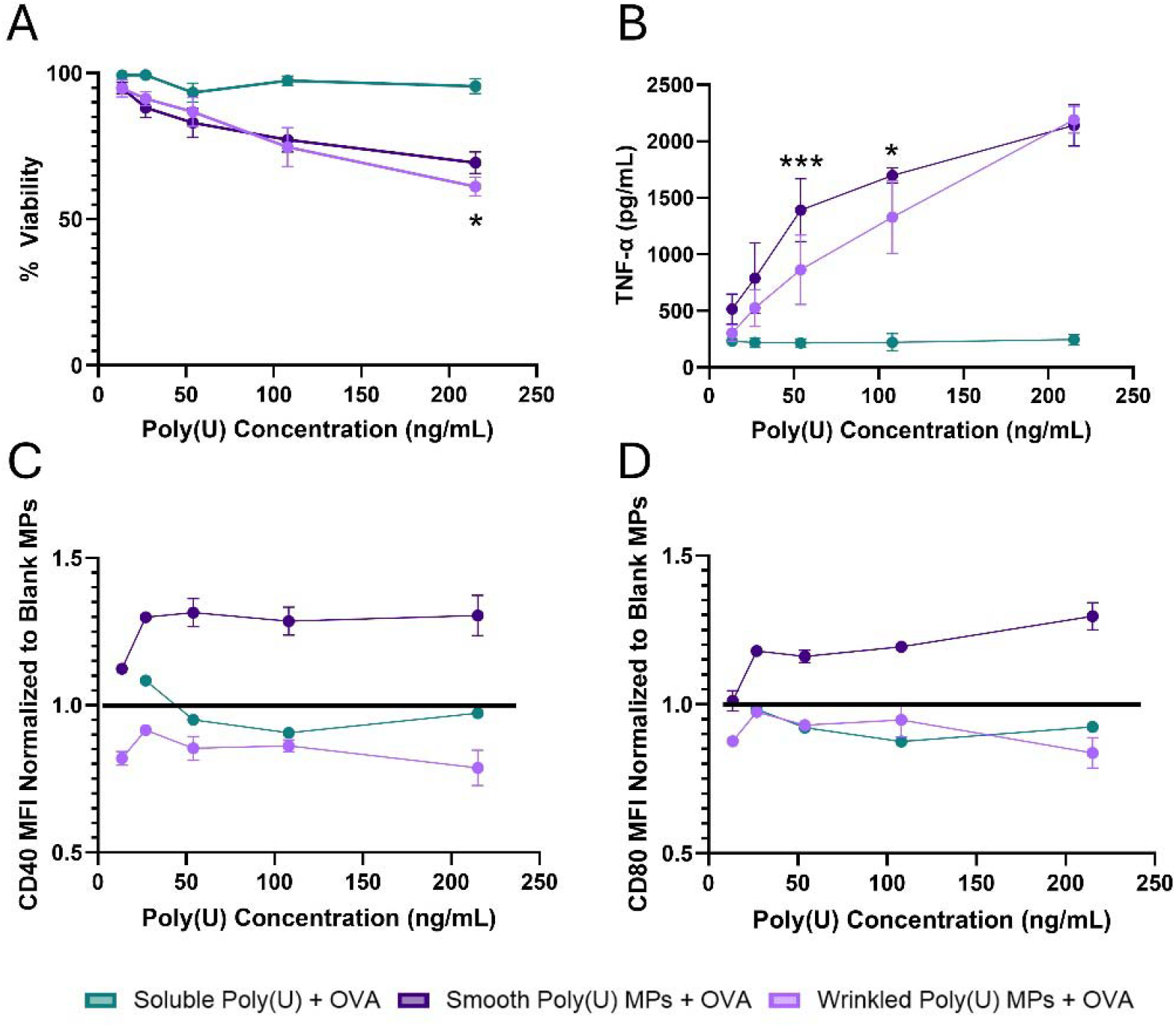
Smooth poly(U) MPs yielded the greatest DC2.4 activation. DC2.4s were incubated with wrinkled or smooth poly(U) MPs and soluble poly(U) for 24 hours. After 24 hours (A) cell viability was measured by a lactose dehydrogenase (LDH) assay, (B)TNF-α concentration in the culture supernatant was measured by cytokine ELISA, (C) CD40 MFI normalized to Blank MPs, and (D) CD80 MFI normalized to Blank MPs were measured by flow cytometry. Data presented as mean ± standard deviation.^*^ p≤0.05^***^ p ≤ 0.001. State what the “*” is significant to, soluble or bn formulations.

Poly(U) is a TLR7/8 agonist which leads to overall DC activation and cytokine responses such as TNF-α and IL-6. To understand how MP morphology affects DC activation, TNF-α and IL-6 were measured in the supernatant of cells treated for 24 hours. Both smooth poly(U) MPs and wrinkled poly(U) MPs induced TNF-α cytokine production in a dose dependent manner **(Fig 2B)** that surpassed soluble poly(U) and blank controls **(Fig 2B, Fig S3B)**. Smooth poly(U) MPs had significantly higher TNF-α production at 215 ng/ml concentration, which is critical as this concentration as it had no significant viability differences from controls **(Fig 2A, Fig S3B)**. Wrinkled poly(U) had high IL-6 production at the highest concentration of 215 ng/mL, however at this concentration the wrinkled poly(U) MPs had significant viability loss which may account for the cytokine production **(Fig S3C, Fig 2A)**.

The DCs were also assessed for immune activation and costimulatory markers via flow cytometry. Previous studies have shown that elevated TNF-α levels are linked to increased expression of CD40 and CD80, key co-stimulatory molecules involved in antigen presentation and the activation of T cells, which are essential for mounting effective immune responses against pathogens [38-40]. Complementing the increase in TNF-α, smooth poly(U) MPs had increased mean fluorescent intensities (MFI) of CD40 **(Fig 2C)** and CD80 **(Fig 2D)** when normalized to Blank MP controls. Wrinkled poly(U) MPs did not influence these DC costimulatory markers, despite having some TNF-α production. These increases in CD40, CD80, and production of TNF-α without significant viability loss for smooth poly(U) MPs affirm their potential as an adjuvant system.

Given the ability to stimulate an innate immune response in DCs, mice were vaccinated with a PBS control without OVA, or soluble poly(U), blank MPs, smooth poly(U) MPs, or wrinkled poly(U) MPs all with 10 mg of OVA. Animals in the treatment groups received a 10 mg soluble OVE protein dose mixed with the MPs as we have previously shown that absorption of OVA protein on the MP surface is as adequate for vaccination [41]. A subcutaneous injection route was chosen to mimic vaccines currently on the market including measles, mumps, and rubella (MMR), chickenpox, and diphtheria, tetanus, acellular pertussis (DTaP) [42]. Vaccinations were given on a prime (day 0), boost (day 21), boost (day 35), schedule; animals were sacrificed ten days following the final vaccination (day 45). B and T cell responses were then assessed in the draining inguinal lymph node (iLN) and spleen. This is an important timepoint as we have previously shown that by day 10 after final vaccination, a majority of Ace-DEX MPs have left the subcutaneous site and drained to the iLN, thus priming the immune system [34, 43]. Cells isolated from the iLNs and splenocytes were untreated or stimulated with OVA protein, MHC-I OVA_257–264_ (SIINFEKL) peptide, or MHC-II OVA_323–339_ (ISQAVHAAHAEINEAGR) peptide. MHC-I and MHC-II stimulation was used to gauge cytotoxic T-cell and helper T-cell responses, respectively, with OVA also stimulating MHC-II.

We observed differential responses in the iLNs across groups stimulated with OVA protein. Vaccination with either smooth and wrinkled poly(U) MPs with OVA had similar frequencies of CD4 T cells to the PBS control **(Fig 3A)**. Wrinkled MPs had a slight trend towards increased CD8 T cell frequency, although it was not significant **(Fig 3B)**. Vaccination with smooth poly(U) MPs with OVA had increased B cell frequency compared to PBS and soluble poly(U) with OVA controls **(Fig 3C)**. Yet, blank MPs with OVA had the most striking response, where their CD4 T cell frequency was non-statistically lowered **(Fig 3A)** but their B cell **(Fig 3C)** and GC B cell frequencies **(Fig 3D)** were significantly increased compared to all other groups. This indicates that blank MPs with OVA significantly biased the immune response to a B cell response locally in the draining iLN, perhaps at the expense of the CD4 T cell response, while smooth and wrinkled poly(U) MPs appear to have more balanced T and B cell responses. Similar observations were made when assessing total iLN cell counts **(Fig S4A)**. We did not note any significant differences in central versus effector memory T cells in the CD4 or CD8 compartments. LNs were also stimulated with OVA’s MHCI and MHCII peptides **(Fig S4B-E)**. We continued to observe that Blank MPs strongly skewed the immune response towards a B cell response.

**Figure 3.**
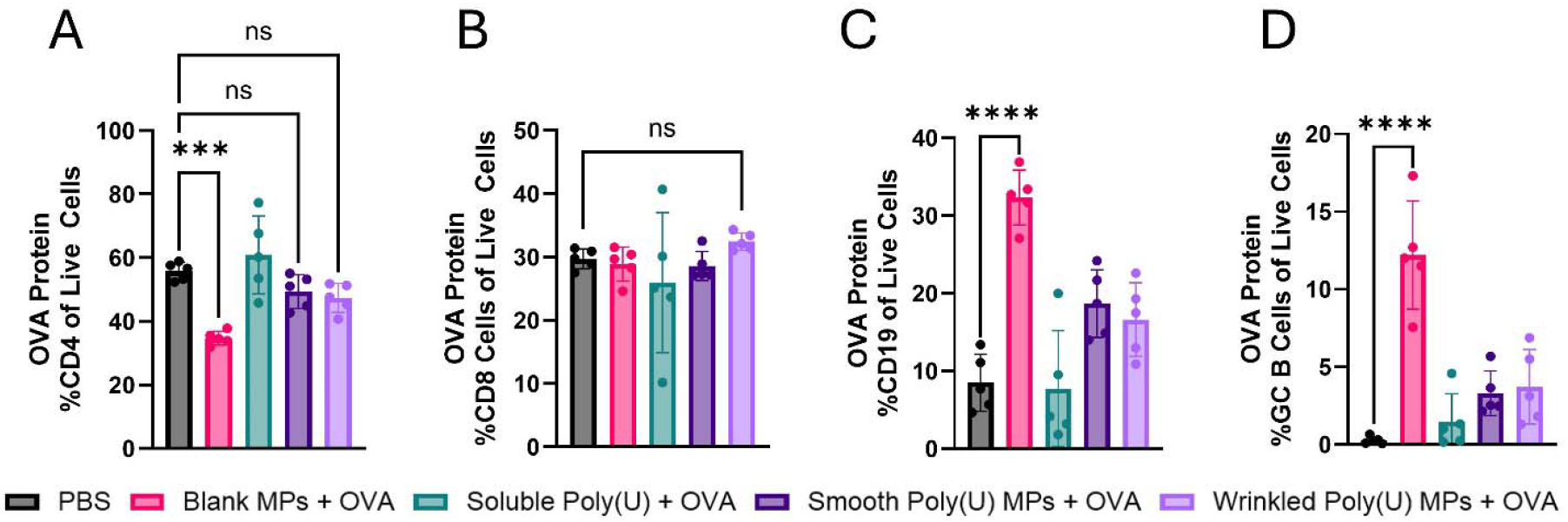
After vaccination, blank MPs with OVA bias cells towards a local B cell response. Using a prime-boost-boost vaccination schedule, inguinal lymph nodes were collected on day 45 and restimulated with full OVA protein and analyzed on flow cytometry. **(A)** Percentage of CD4 T cells, **(B)** percentage of CD8 T cells, **(C)** percentage of CD19 B cells, and **(D)** percentage of germinal center (GC) B cells. Data presented as mean ± standard deviation. ^*^ p≤0.05, ^***^ p ≤ 0.001, ^****^ p ≤ 0.001.

Spleens were also assessed from vaccinated mice and restimulated with OVA and its MHC-I, and MHC-II peptides, which had vastly different responses from the lymphocytes. At baseline, wrinkled poly(U) MPs trended towards greater frequency of CD19+ B Cells than all other groups irrespective of stimulant **(Fig 4A)**, indicating that B cells have systemically been primed from the PBS baseline. This trend was consistent across all stimulations with OVA as well **(Fig 4B-D)**. Assessing GC B cells, wrinkled poly(U) MPs had increased GC B cells out of all CD19 B cells compared to PBS, Blank MPs, and soluble poly(U**(Fig 4 E-H)**,). When restimulated with OVA and its MHC-II peptide, smooth poly(U) MPs had more GC B cells out of CD19 B cells than wrinkled poly(U) MPs. However, since wrinkled poly(U) MPs had a greater frequency of B cells at baseline, we concluded that this group had the greatest B cell response systemically.

**Figure 4.**
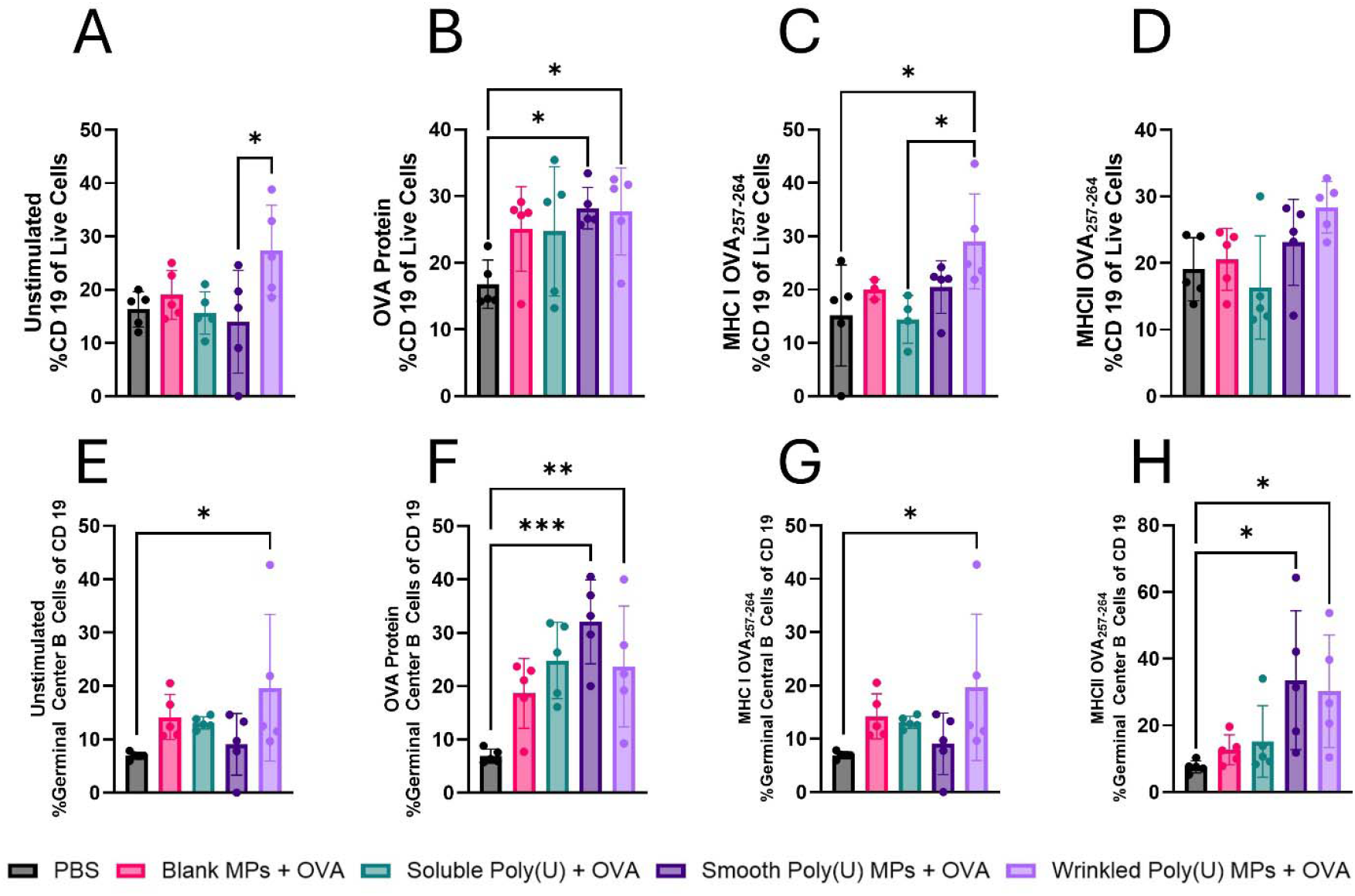
After vaccination, wrinkled poly(U) MPs with OVA bias cells towards a systemic B cell response. On day 45 after the first vaccination, (A-H) splenocytes were isolated from mice, made into single-cell suspensions (n = 5), and stimulated with (A,E) nothing, (B,F) OVA, (C,G) MHC-I, or (D,H) MHC-II and to measure for (A-D) B Cells (CD19), and (E-H) Germinal Center B Cells (GL7+, CD38+). Data from two-way ANOVA is shown as mean ± SD. ^*^ p≤0.05, ^**^ p≤0.01, ^***^ p≤0.001.

**Figure 5.**
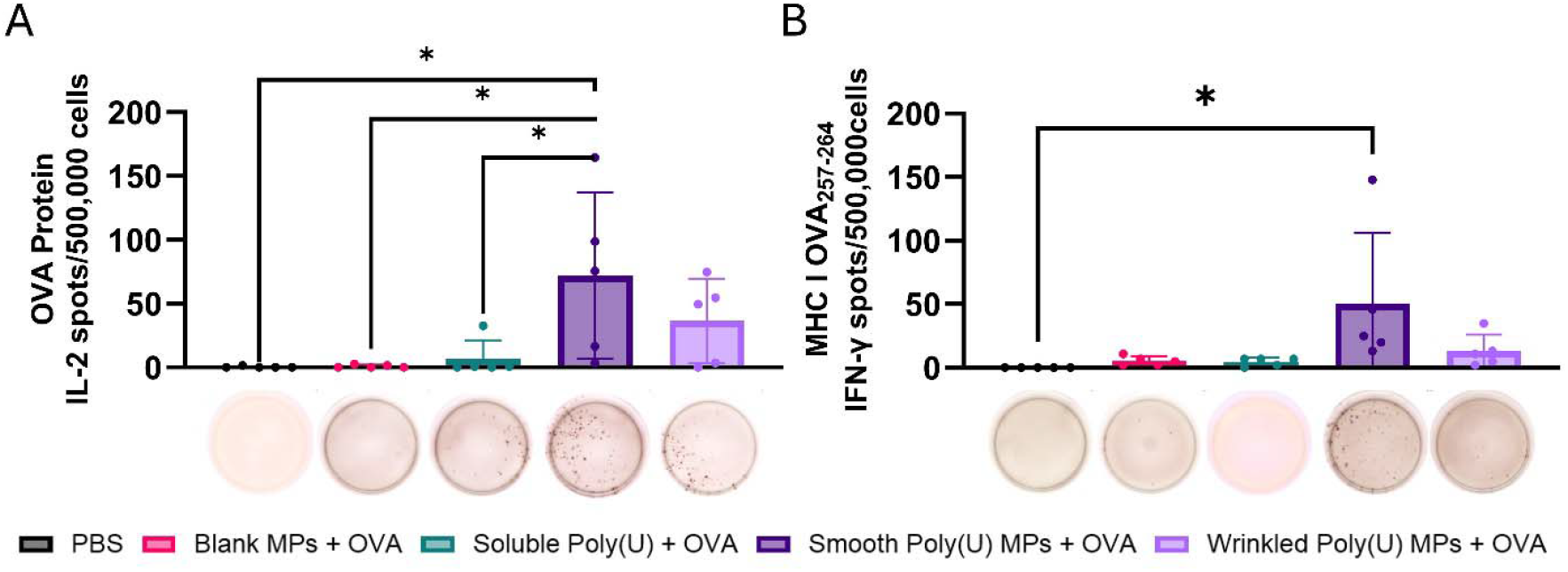
Smooth poly(U) MPs have a strong recall immune response during antigen recall by splenocytes from day 42 vaccinated mice. Spleens were isolated, made into single-cell suspensions to evaluate their cellular response on day 45 post prime + boost + boost vaccination, and stimulated and had high expression of smooth poly(U) MPs from ELISpot for (A) OVA and IL-2 and (B) MHC-I peptide and IFN-γ. Representative images of an ELISpot well are shown under their corresponding vaccination group; data is from two-way ANOVA and shown as mean ± standard deviation. ^*^ p ≤ 0.05.

To assess the T-cell response, we did ELISpot for cytokines. An increase in IFN-γ and IL-2 production was observed in splenocytes from mice vaccinated with smooth poly(U) MPs in a statistically significant response to both OVA and MHC-I OVA peptide stimulation. An MHC-I response is indicative of a cytotoxic T-cell response. IL-2 and IFN-γ are primarily produced by T cells and indicate, generally, proliferation at high concentrations and activation at most concentrations, respectively. IFN-γ production is important when considering vaccine development for a variety of disease states because it promotes T cell responses [44]. Boosting T cell responses in a vaccine would contribute to developing treatments for cancer [45, 46], bacterial infections [47, 48], and viral infections [49]. While there was a cytokine response to restimulation with OVA peptide and MHC-I immunodominant peptide there were not any cytokine responses to restimulation with MHC-II immunodominant peptide **(Fig S5A-B)**. The IL-2 and IFN-γ response to OVA and MHC-I immunodominant peptide indicates activation of antigen-specific T cells, highlighting smooth poly(U) MPs’ ability to trigger a cytotoxic T cell response [50, 51].

## Conclusions

To enhance the delivery of to its intracellular receptor, poly(U) was formulated in an Ace-DEX MP via spray drying. On average, wrinkled MPs had a greater diameter than smooth MPs. Both smooth and wrinkled MPs had similar release kinetics (**Fig 1A-C**). In DCs in vitro, smooth poly(U) MPs were able to elicit an increased cytokine response and more co-stimulation than wrinkled poly(U) MPs (**Fig 2)**. Both smooth and wrinkled MPs cause higher cytokine production than soluble poly(U). Results from in vivo studies show Blank MPs biased the immune response to a B cell response locally in LNs whereas smooth and wrinkled poly(U) MPs appeared to have a more balanced B and T cell response **(Fig 3)**. In splenocytes of mice treated with MPs, wrinkled poly(U) MPs were the most effective in increasing B cell frequency **(Fig 4)**. Whereas the in vitro stimulation found that smooth poly(U) MPs were more adept at producing a cytokine response **(Fig 2B)**, in vivo, wrinkled poly(U) influenced B-cell frequency **(Fig 4A-4C)**. Overall, we show that wrinkled poly(U) MPs caused the highest frequency of B cells across splenocytes and iLNs. In future studies, a combined wrinkled and smooth formulation should be investigated for a balanced response between B cells and T cells.

## Credit Authorship Contribution Statement

**Sophia Ly:** Writing – review & editing, Writing – original draft, Methodology, Formal analysis, Data curation. **Nicole Rose Lukesh:** Writing – review & editing, Writing – original draft, Methodology, Formal analysis, Data curation. **Grace Williamson, Connor Murphy, Sophie Mendell, Ryan Woodring, Kiersten Clark** and **Alexandra Lopez:** Data Curation. **Eric M. Bachelder:** Writing – review & editing, Writing – original draft, Supervision, Resources, Methodology. **Kristy M. Ainslie:** Writing – review & editing, Writing – original draft, Supervision, Resources, Methodology, Funding acquisition.

## Declaration of Competing Interest

The authors declare the following financial interests/personal relationships which may be considered as potential competing interests: Kristy Ainslie reports financial support was provided by National Institute of Health. Kristy Ainslie reports a relationship with National Institutes of Health that includes: funding grants. None If there are other authors, they declare that they have no known competing financial interests or personal relationships that could have appeared to influence the work reported in this paper.

## Acknowledgements

This work was supported by the internal funds at the University of North Carolina – Chapel Hill and the National Institutes of Health. The authors would also like to give thanks to the Lazear lab at UNC for the use of their ELISpot reader. The work was also performed with the Chapel Hill Analytical and Nanofabrication Laboratory, CHANL, a member of the North Carolina Research Triangle Nanotechnology Network, RTNN, which is supported by the National Science Foundation, Grant ECCS-1542015, as part of the National nanotechnology Coordinated Infrastructure, NNCI. Flow cytometry was performed at the UNC Flow Cytometry Core Facility (RRID:SCR_019170) which is supported in part by P30 CA016086 Cancer Center Core Support Grant to the UNC Lineberger Comprehensive Cancer Center, North Carolina Biotech Center Institutional Support Grant 2017-IDG-1025, and by the National Institutes of Health 1UM2AI30836-01. Graphical figures were produced in Biorender (biorender.com) and data figures were produced in GraphPad Prism (graphpad.com).

## Notes

### Competing Interest Statement

The authors have declared no competing interest.

