## Supplemental Data for "Controlled Release of Poly(U) via Acetalated Dextran Microparticles for Enhanced Vaccine Adjuvant Delivery"

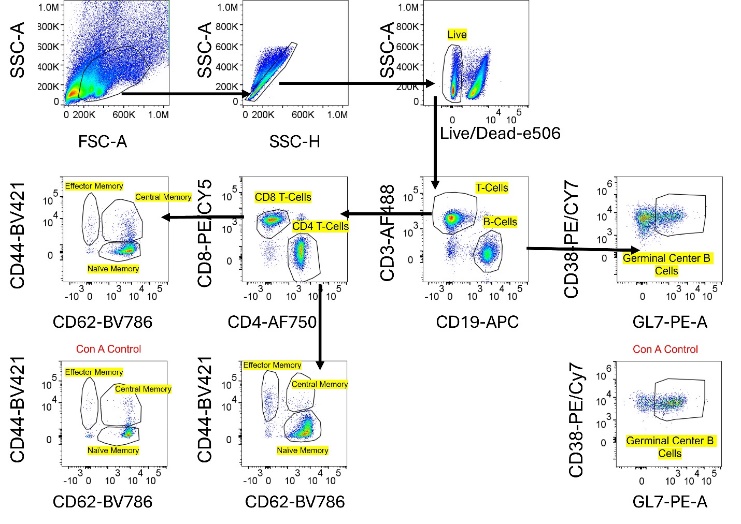

Supplemental Figure 1. Flow Gating Strategy for Splenocytes

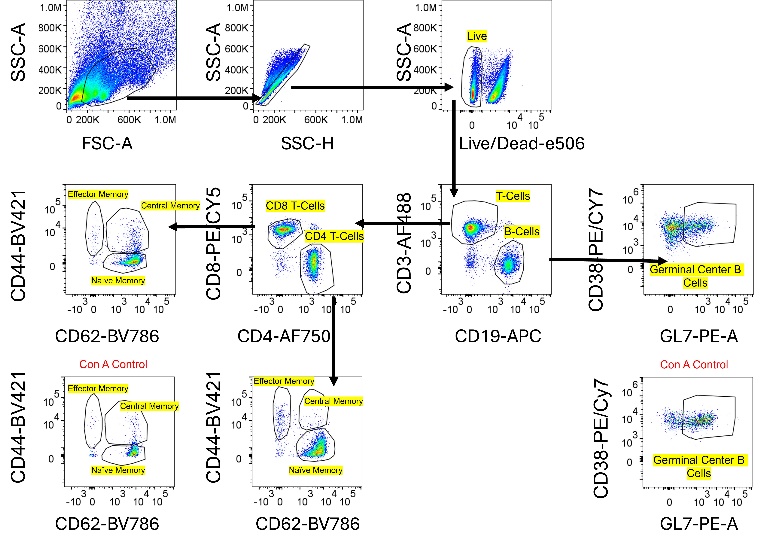

Supplemental Figure 2. Flow Gating Strategy for Lymphocytes

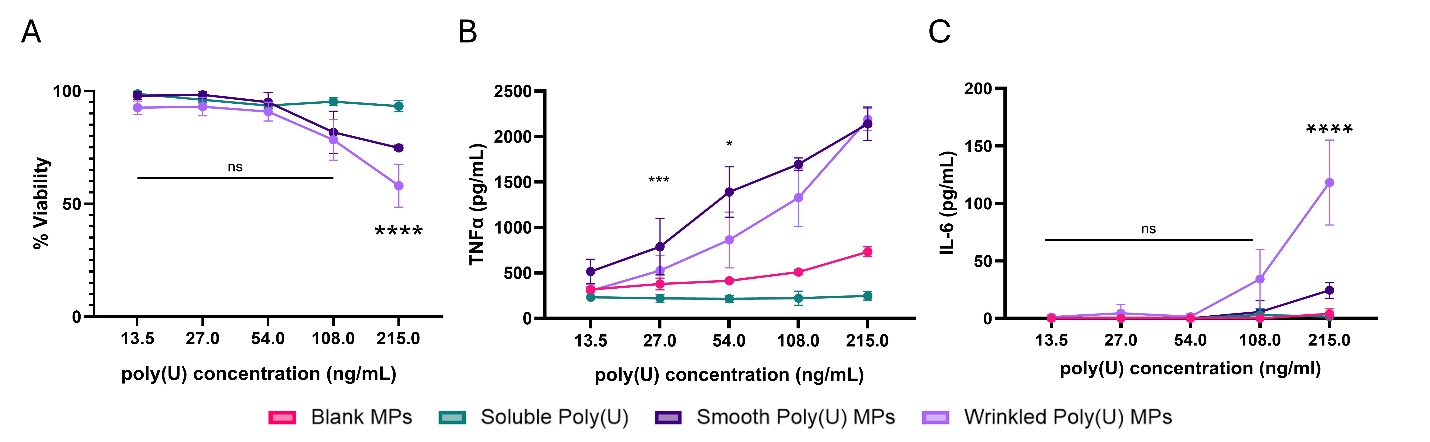

Supplemental Figure 3. Wrinkled poly(U) MPs had lower percent viability and higher IL-6 activation. DC2.4s were incubated with Wrinkled or smooth poly(U) MPs and soluble poly(U) for 24 hours. After 24 hours (A) cell viability was measured with Cell Titer Blue assay and (B) TNF-α IL-6 concentration in the culture supernatant was measured by cytokine ELISA. **** p ≤ 0.001.

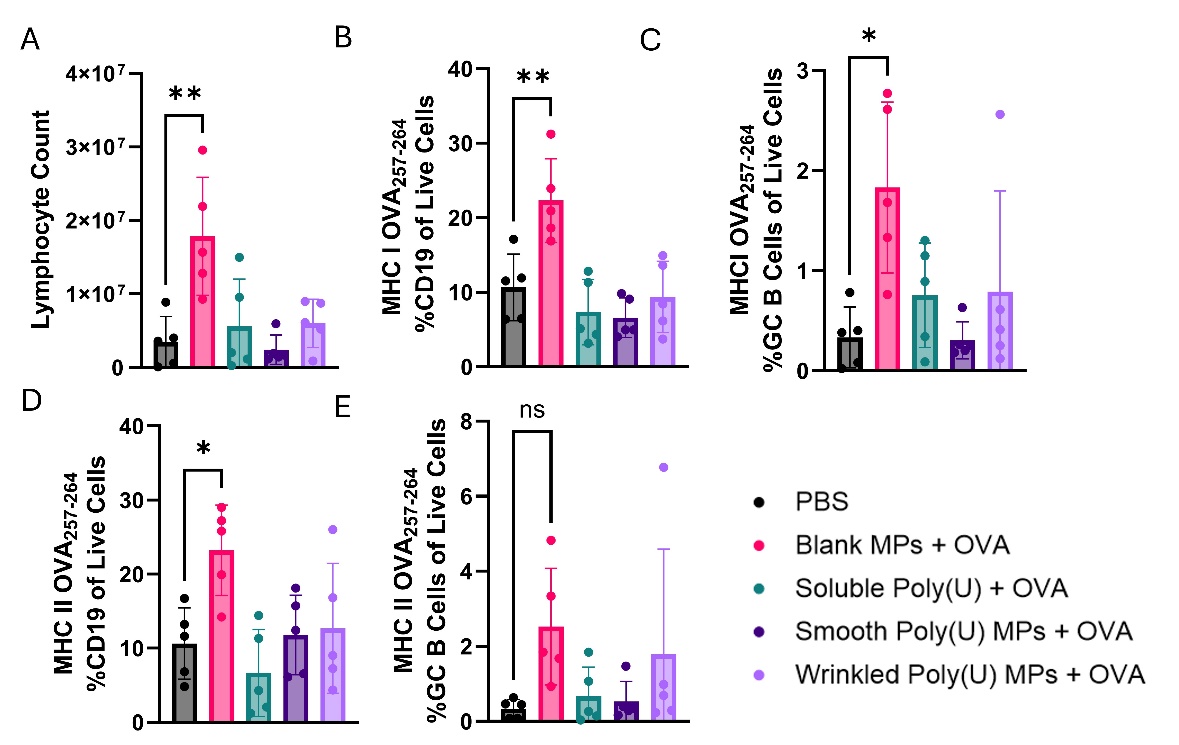

Supplemental Figure 4. After vaccination, Blank MPs with OVA bias cells towards a local B cell response. Using a prime-boost-boost vaccination schedule, inguinal lymph nodes (iLNs) were collected on day 45 and restimulated with full MHC-I and MHC-II peptide and run on flow cytometry. **(A)** Lymphocyte Count **(B)** percentage of CD19 in MHC-I restimulation, **(C)** Percentage of GC B cells in MHC-I restimulation, **(D)** percentage of CD19 in MHC-II restimulation and **(E)** percentage of germinal center (GC) B cells in MHC-II restimulation. Data presented as mean ± standard deviation. * p≤0.05 ** p≤0.01

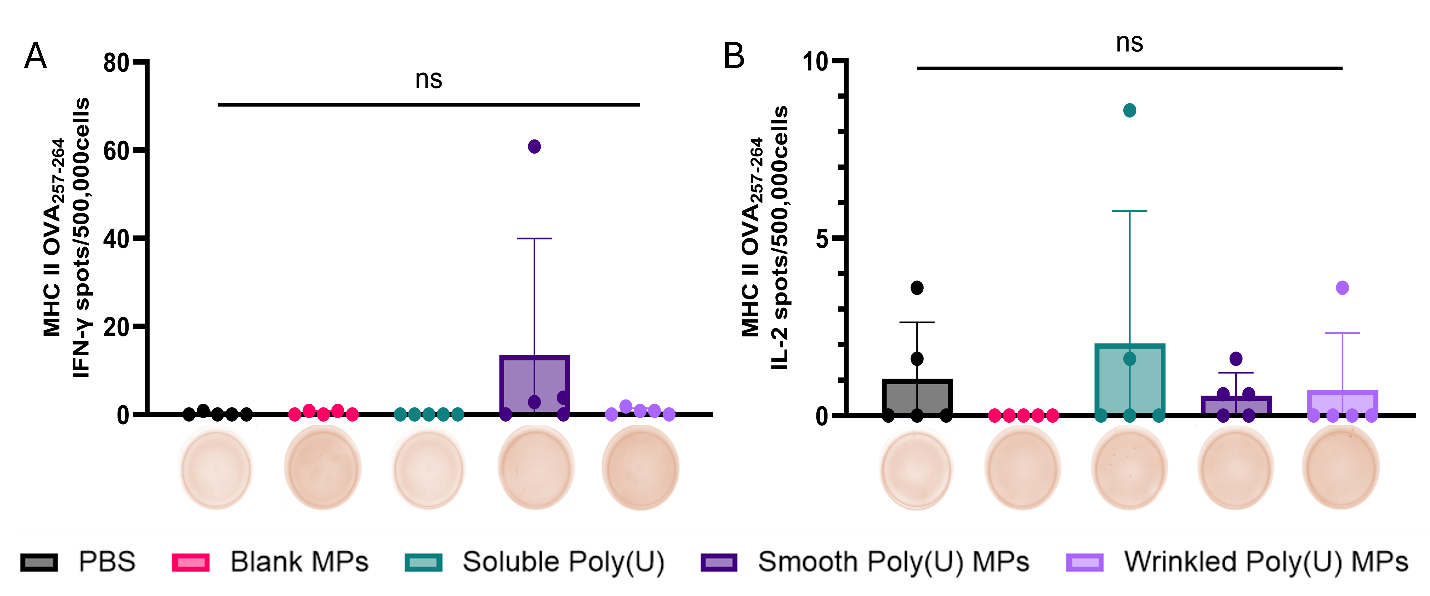

Supplemental Figure 5. No treatment groups have a strong recall immune response during antigen recall by splenocytes from day 42 vaccinated mice when restimulated with MHC-II. Spleens were isolated, made into single-cell suspensions to evaluate their cellular response on day 45 post prime + boost + boost vaccination, and stimulated and had no significant expression from ELISpot for (A) MHC-II and IFN-γ and (B) MHC-II peptide and IL-2. Representative images of an ELISpot well are shown under their corresponding vaccination group; data is from two-way ANOVA and shown as mean ± standard deviation.
